# Genomic relationships reveal significant dominance effects for growth in hybrid *Eucalyptus*

**DOI:** 10.1101/178160

**Authors:** Biyue Tan, Dario Grattapaglia, Harry X. Wu, Pär K. Ingvarsson

## Abstract

Non-additive genetic effects can be effectively exploited in control-pollinated families with the availability of genome-wide markers. We used 41,304 SNP markers and compared pedigree *vs.* marker-based genetic models by analysing height, diameter, basic density and pulp yield for 338 *Eucalyptus urophylla* x *E.grandis* control-pollinated families represented by 949 informative individuals. We evaluated models accounting for additive, dominance, and first-order epistatic interactions (additive by additive, dominance by dominance, and additive by dominance). We showed that the models can capture a large proportion of the genetic variance from dominance and epistasis for growth traits as those components are typically not independent. We also show that we could partition genetic variances more precisely when using relationship matrices derived from markers compared to using only pedigree information. In addition, phenotypic prediction accuracies were only slightly increased by including dominance effects for growth traits since estimates of non-additive variances yielded rather high standard errors. This novel result improves our current understanding of the architecture of quantitative traits and recommends accounting for dominance variance when developing genomic selection strategies in hybrid *Eucalyptus*.

## 1. Introduction

Hybrids between inbred lines within species or between different species are commonly used for commercial production in both crops and tree species. The main reason of conducting crosses between pure lines of a single species or between contrasting species is the exploitation of hybrid superiority (heterosis) or to combine complementary traits of different species [1-3]. The major goal of such hybrid breeding programs is to identify the best performing hybrid individuals for subsequent cultivar development [4]. Moreover, the best performing individuals of the contrasting populations can be used as parents of a new breeding population in further long-term breeding strategies [5, 6]. In forest trees, the worldwide production of hybrid poplar and eucalyptus are two successful examples of hybrid breeding [7].

Our current understanding of the occurrence of heterosis is based on genetic theory of dominance effects [8] which has subsequently been extended to include all non-additive genetic effects (dominance and epistasis, [9]). Dominance arises due to interactions between alleles at the same locus whereas epistasis is due to interactions between alleles at different loci [10]. While some studies have found that dominance variance can contribute substantially to trait variation in forest trees [11], others have shown very little contribution of dominance [12, 13]. The importance of non-additive genetic variance relative to additive genetic variance also changes across different ages when a trait is measured [14]. Overall, there have been only a few reliable estimates of non-additive genetic parameters in forest tree species. Genetic variance and broad sense heritability (H^2^) are expected to be higher than the corresponding additive variance and narrow-sense heritability (h^2^) if there is significant non-additive genetic variance and the 
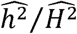
 ratios reported for traits in forest trees have ranged from 0.18 to 0.84 (
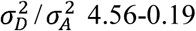
) [7, 15, 16]. For *Eucalyptus* hybrids, the relative contribution of dominance has been shown to vary between traits and species combinations. It has been reported that rooting ability, flowering time, drought and freezing resistance were all inherited in a predominantly additive manner (reviewed in [17]), while partial dominance was detected for freezing resistance in F_1_ hybrids of *E. camaldulensis* × *E. globulus* and *E. torelliana* × *E. citriodora*, respectively [18]. Dominance effects seem to be important and widespread for growth traits [1, 19-21] and a ratio of dominance to additive variance close to 1.2 was estimated during the growth period for the *E. grandis x E. urophylla* hybrid [11]. On the other hand, previous reports have indicated that wood density is inherited in an additive manner in virtually all *Eucalyptus* species combinations examined to date ([22], reviewed in [17]). Finally, pulp yield appears to show dominance or partial dominance towards the low yielding parents [18].

Although many studies have estimated non-additive effects, it is challenging to obtain accurate estimates for non-additive genetic variances using pedigree information for a number of reasons. First, large full-sib families or deep pedigree trials with vegetatively propagated populations (clonal trials) are required to accurately estimate non-additive effects [10]. Second, non-additive genetic effects could be confounded with species, provenance and/or environmental effects [23-27]. An additional limitation is imposed by the potential uncertainty of the pedigree information, which may contain parentage errors such that estimates are based on the expected and not the realized degree of genetic relationship. This can be particularly problematic for forest trees where controlled crosses are laborious and prone to errors or pollen contamination.

Recent advances of high-throughput genotyping technologies and the availability of whole genome single nucleotide polymorphism (SNP) marker panels have made it feasible to estimate genetic variance components based on genomic data using, for example, realized genomic relationships (GBLUP) [28]. Additive, dominance and epistasis variance components can then be estimated by constructing genome-wide SNP marker-based relationship matrices that allow more precise separation of confounding factors compared to estimation of genetic variance based on pedigrees [29, 30]. Most initial GBLUP studies in forest trees focused solely on estimating additive genetic variances [31-40] However, a few recent studies have also reported the contribution of non-additive effects to phenotypes [41-44]. Analysis of simulated data indicate that including dominance could result in higher genetic gains in crossbred population [45] and adding dominance effects can increase the prediction accuracy of phenotype when non-additive variation constitute a considerable proportion of the phenotypic variance [44, 46]. Results for prediction of genetic values have been contradictory, however. For example, Muñoz et al. [29] found that there was little improvement in prediction accuracy of phenotypic values for height in loblolly pine when accounting for non-additive variation. Similar results have also been found in hybrid *Eucalyptus* populations. For example, although a large dominance variance component was found for height, it led to a very small improvement in predicting phenotypic values [41,47]. Due to the conflicting results regarding the relative importance of non-additive effects in predicting trait values and potentially selecting candidates with best genetic performance, the objectives of this study were to compare the performance of pedigree-based and genomics-based models including both additive and non-additive effects in a hybrid *Eucalyptus* population. Because we previously identified inconsistences between pedigree-based and realized relationships [48], we reconstruct the 'true’ pedigree using genotype information. We focused on growth traits at age 3 and 6 years and wood property traits and assessed the impact of including non-additive effects on the predictive ability. i.e. the correlation between genetic values and phenotypes, of the various models employed.

## 2. Materials and methods

### 2.1. Outcrossed Eucalyptus progeny test, phenotype data and genotyping

The progeny population and their phenotypic and genotypic data used in this study have been previously described in Tan *et al.* [48]. Briefly, the progeny test was established by controlled crossing of 86 *E. urophylla* and 95 *E. grandis* trees resulting in 476 full-sib families with 35 individuals per family, and the field test was grown in a randomized complete block design with single-tree plots and 35 blocks in the trial. The present study is based on a subset of this trial, involving 958 individuals from 338 full-sib families after removing outlier trees likely due to selfing or general health issues. The number of individuals in each full-sib family ranged from one to 13 with the median of 2.44. Height and circumference at breast height (CBH) were measured at age three and six years and wood basic density and pulp yield were determined using Near-Infrared Reflectance spectra at the age of five years. All 958 trees were genotyped using the Illumina Infinium EuCHIP60K that contains probes for 60,904 SNPs [49]. After quality-control based on greater than 70% call rates of both SNPs and samples, minor allele frequencies greater than 0.01 and Hardy-Weinberg equilibrium (p-value < 1×10^−6^), 41,304 SNPs were retained for 949 samples. SNPs with less than 2.1% missing information were imputed by BEAGLE 4.0 and used in all subsequent analyses [48].

### 2.2. Pedigree reconstruction

Since we found considerable inconsistencies between the registered pedigree and the realized relationships in our previous study [48], we carried out a parentage assignment test in this study to better understand the reasons of these inconsistencies and to construct a pseudo-pedigree that was later used to estimate genetic parameters and make predictions compared to genomic-based ones. We assigned parentage to all 949 progenies using the program SNPPIT [50], which employs SNP markers to identify the most likely parent pairs for all progenies based on a pool comprising 90 *E. grandis* and 84 *E. urophylla* parental candidates. The program uses a likelihood-based categorical assignment method and a Monte Carlo simulation to assess confidence of parentage assignments based on false discovery rate (FDR) calculations. We only accepted assignments where the estimated FDR was less than 5%. We repeated the SNPPIT analyses 100 times by randomly sampling 96 independent SNPs without repetition as suggested by Anderson [50] and assumed a SNP genotyping error rate of 1% for each run. Before we ran SNPPIT, 10,213 independent SNPs were obtained from PLINK through LD-pruning (r^2^ < 0.2) [51]. In addition, we found that all parents were not independent of each other and a few parents displayed relatedness up to 0.7, suggesting a relationship greater than full-sibs. For this reason, we summarized the frequencies of assigned parents after 100 repetitions and selected those that were assigned as pseudo-parent(s) candidates with greater than 50% frequency for each of the 949 progeny individuals.

### 2.3. Phenotypic trait adjustments

Prior to the analyses of additive and non-additive effects, phenotypic traits were adjusted for environmental variation by fitting the following linear mixed model to the phenotypic data:

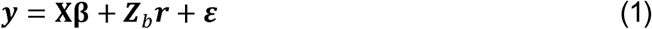

where *y* is the vector of phenotypic observation, **β** is the vector of fixed effects (overall mean), ***r*** is the vector of random block effects following 
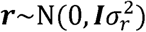
, where 
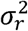
 is the block variance, and ***Z*_b_** is block design matrix, *ε* the vector of random residual. The residual R matrix is structured as

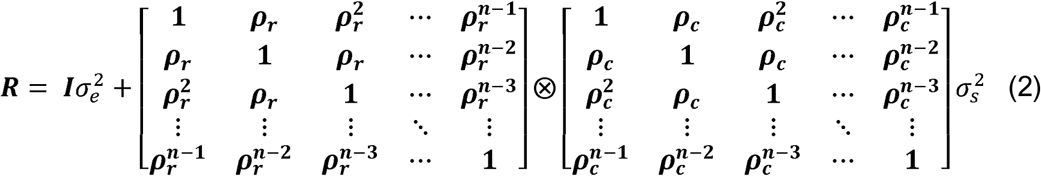

where ⊗ represent the Kronecker product [52], ***ρ****_r_* and ***ρ****_c_* are the autoregressive first order correlations in the row and column directions, respectively. Model parameter estimation for Equation 1 was carried out using a residual maximum likelihood (REML) method as implemented in ASReml 4.1 [53]. Finally, adjusted phenotypes of each trait were obtained by subtracting effects of random block and spatial position. These adjusted phenotypes were used for all further analyses in the study.

### 2.4. Pedigree and genomic relationship matrices

The pedigree co-ancestry coefficients were estimated based on the pedigree of the female and male parent population. The diagonal elements (*i*) of the additive relationship matrix (**A**) were calculated as 
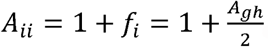
, where *g* and *h* are the *i*'s parents; while the off-diagonal element is the relationship between individual *i*th and *j*th calculated as 
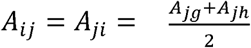
 [10]. The off-diagonal elements between individual *i*th and *j*th in the dominance relationship matrix (**D**) can be computed as 
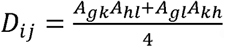
, where *g* and *h* are the *i*'s parents and *k* and *l* are the *j*'s parents; whereas the diagonal elements are all *D_ii_* = 1.[10]. Both **A** and **D** relationship matrices were calculated using the “kin” function from the “synbreed” package in R [54].

The genomic-based additive relationship matrix was estimated using the formula developed by VanRaden [55]: 
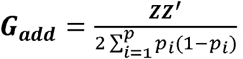
, where ***Z*** is a mean-centred matrix of *n*individuals by *m* SNPs following ***M – P, M*** is the genotype matrix coded as 0, 1 and 2 according to the number of alternative alleles, and ***P*** is the matrix of average locus scores *2p_i_*, where *p_i_* is the *i*th allele frequency and 
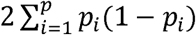
 is the variance of markers summed cross all loci. The genomic-based dominance relationship matrix was estimated as 
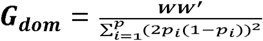
, where ***W*** is the matrix containing –2(1 − *p_i_*)^2^ for the alternative homozygote, *2p_i_* (1 – *p_i_*) for the heterozygote, and 
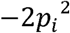
 for the reference allele homozygote of *i*th SNP [56].

The relationship matrices due to the first-order epistatic interactions were computed using the Hadamard product (element by element multiplication, denoted #). Under the pedigree-based relationship matrix, additive × additive terms ***E_AA_*** = A#A, additive × dominance terms ***E_AD_*** = A#D, and dominance × dominance terms ***E_DD_*** = D#D; while under the genomic based relationship matrices, additive × additive terms ***G_AA_*** = ***G_add_#G_add_***, additive × dominance terms ***G_DD_*** = ***G_dom_#G_dom_***[57].

### 2.5. Variance components and heritability models

Estimates of variance components for each trait were obtained using the best linear unbiased prediction (BLUP) method in three univariate models that included either only additive (A), additive and dominance (AD), or additive, dominance and epistatic (ADE) genetic effects as follows:

For the model with additive effects only (A):

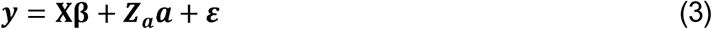

where ***y*** is the vector of adjusted phenotypes after elimination of environmental effects, **β** is the vector of fixed effects (overall mean), and ***ε*** is a vector of the random residual effects following 
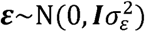
, where 
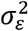
 is the residual variance. ***a*** is the vector of additive genetic effects, which following 
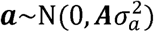
 for pedigree-based relationship matrix, where ***A*** is the additive numerator relationship matrix as described above and 
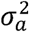
 is the corresponding additive genetic variance. When using the genomic-based relationship matrix for the analyses, ***A*** was substituted with ***G_add_*** and ***a*** yielding 
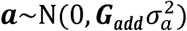
, where ***G_add_*** is the marker-based relationship matrix as de s c ribed above (Table 1). ***X*** and ***Z_a_*** are incid e nce matrices relating fixed and random effects to measurements in vector ***y***.

The extended model including dominance terms (AD) was:

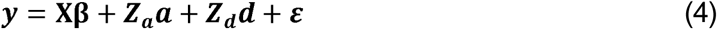

where *d* is the vector of the random dominance effect following 
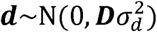
 for the variance components analysis using pedigree-based relationship matrix, where *D* is the dominance numerator relationship matrix as mentioned above and 
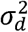
 is the corresponding dominance genetic variance. For analysing dominance genetic variance components using the genomic-based relationship matrix, ***D*** was replaced by ***G_dom_***(Table 1). Other parameters are as described above.

**Table 1.** Additive and non-additive genetic models and the associated relationship matrices.

| Matrix type | Model type | Fixed effect | Relationship matrices related to the model |  |  | Code |
| --- | --- | --- | --- | --- | --- | --- |
|  |  |  | Additive | Dominance | Epistasis |  |
| Registered (expected) pedigree ($P_{\text{exp}}$) | A | Mean | $A$ | | | $P_{\text{exp}}A$ |
| | AD | Mean | $A$ | $D$ | | $P_{\text{exp}}AD$ |
| | ADE | Mean | $A$ | $D$ | $E_{AA}, E_{AD}, E_{DD}$ | $P_{\text{exp}}ADE$ |
| Pseudo-pedigree (P) (SNP-estimated parentage) | A | Mean + genetic groups | $A$ | | | PA |
| | AD | Mean + genetic groups | $A$ | $D$ | | PAD |
| | ADE | Mean + genetic groups | $A$ | $D$ | $E_{AA}, E_{AD}, E_{DD}$ | PADE |
| Genotypes (G) (SNP-based genomic relationship matrix) | A | Mean | $G_{add}$ | | | GA |
| | AD | Mean | $G_{add}$ | $G_{dom}$ | | GAD |
| | ADE | Mean | $G_{add}$ | $G_{dom}$ | $G_{AA}, G_{AD}, G_{DD}$ | GADE |

The final model extension including epistatic terms was:

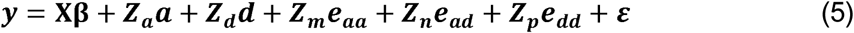

where ***e_aa_*** is the vector of the random additive by additive epistatic effects following 
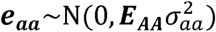
 for the genetic variance components analysis using pedigree-based relationship matrix, ***e_aa_*** is the vector of the random additive × dominance epistatic effects following 
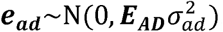
, and similarly, ***e_aa_*** is the vector of the random dominance × dominance epistatic effects following 
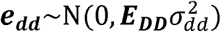
, where 
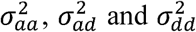
 are the additive × additive, additive × dominance and dominance × dominance epistatic interaction variance, respectively. When we analysed the epistatic interactions using the genomic-based relationship matrix, ***E_AA_***, ***E_AD_*** and ***E_DD_*** matrices were substituted by ***G_AA_***,***G_AD_*** and ***G_DD_***, respectively.

After fitting each model we calculated both narrow-sense and broad-sense heritabilities (h^2^ and H^2^ respectively), which correspond to the proportion of phenotypic variance explained by additive genetic variance only or by additive and non-additive genetic variance combined. Narrow-sense heritability was estimated as 
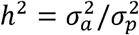
, where 
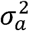
 represented the estimated additive variance and 
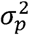
 represented the phenotypic variance which is sum of all the genetic variances and the residual variance. Broad-sense heritability for the A+D model was estimated as 
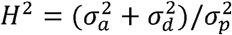
, where 
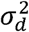
 represented the estimated dominance variance, while H^2^ for the A+D+E model was estimated as 
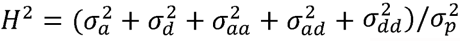
, where 
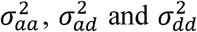
 represented estimated additive × additive, additive × dominance and dominance × dominance epistatic variance, respectively. Finally, we also

^×^calculated the dominance (
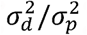
) and epistatic ((
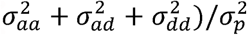
) to phenotypic variance ratios, respectively.

### 2.6. Model comparisons

Models were built by considering different genetic variance compositions and different relationship matrices (Table 1). In this study, we used three relationship matrices, one based on the registered or expected pedigree (P_exp_), one on the SNP-assigned parentage pseudo-pedigree (P) and one built directly from SNP genotypes, i.e. a Genomic Relationship Matrix (G). The models described above were analysed using ASReml 4.1 software [53]. Models were compared using the Akaike Information Criterion (AIC) [58] where AIC was calculated as AIC = 2t - 2ln (*L̂*), where ln (*L̂*) is log-likelihood of the model and the t is the number of variance parameters.

We assessed the precision and dependency among variance components by calculating accumulated eigenvalues of the asymptotic sampling correlation matrix of variance component estimates F, F = *L*^−1/2^ *vL*^−1/2^, where *V* is asymptotic variance-covariance matrix of estimates of variance components and *L* is a matrix containing the diagonal elements of *V* [29]. The eigenvalues were computed using the ‘eigen’ function in R and plots were made relating cumulative percentage of variance explained by the different models with the eigenvalue order.

We evaluated the model fit of the full data set by assessing the correlation between predicted additive genetic values and phenotypes of individuals *r*(*Â_full_*,*Y_full_*) and between predicted total genetic values and phenotypes *r*(*Ĝ_full_*,*Y_full_*)

### 2.7. Models prediction and evaluation

The prediction ability was estimated for all models and relationship matrices. A 10-fold cross-validation scheme with 100 replications was implemented to evaluate the prediction accuracy for different models. For each replication, the dataset was randomly divided into 10 subsets, nine out of the ten partitions were used as the training population to fit a model by using both phenotypes and genotypes while the remaining partition was used as the validation set by removing phenotypic data and then used to predict breeding values or total genetic values for the model in question. The predictive ability of the model was evaluated by estimating the correlation between phenotypes and breeding/genetic values, *r*(*Â_vali_*,*Y_vali_*) or *r*(*Â_vali_*,*Y_vali_*).

## 3 Results

### 3.1. Parentage assignment and pseudo-pedigree creation

In order to compare the results of pedigree-based and genomic-based models, we initially used SNP-based parentage assignment analysis to identify the most likely parents of all progeny individuals since we previously found a large proportion of pedigree errors in the registered pedigree information [48]. Under strict parentage assignment tests, 949 offspring were tested for parentage using the candidate pool of parents. For 850 (89.5%) individuals both parents could be assigned successfully, while for the other 94 (10%) we could only assign a single parent, while for five offspring (0.5%) we could not assign any parent (Figure 1A). For the 944 offspring for which at least one parent was assigned, 72 *E.grandis* and 73 *E.urophylla* were identified parents with range of 2-67 (mean value: 10) crosses per parent. Among these offspring, 207 (21.9%) of their SNP-assigned parents matched the expected parents based on the registered pedigree in the breeders' records. For a set of 586 (62.1%) individuals only the female parent matched the expected one, while for 21 (2.2%) individuals only the male parent matched. For the remaining 130 (13.8%) individuals both the male and female assigned parents did not match the expected ones (Figure 1A).

**Figure 1.**
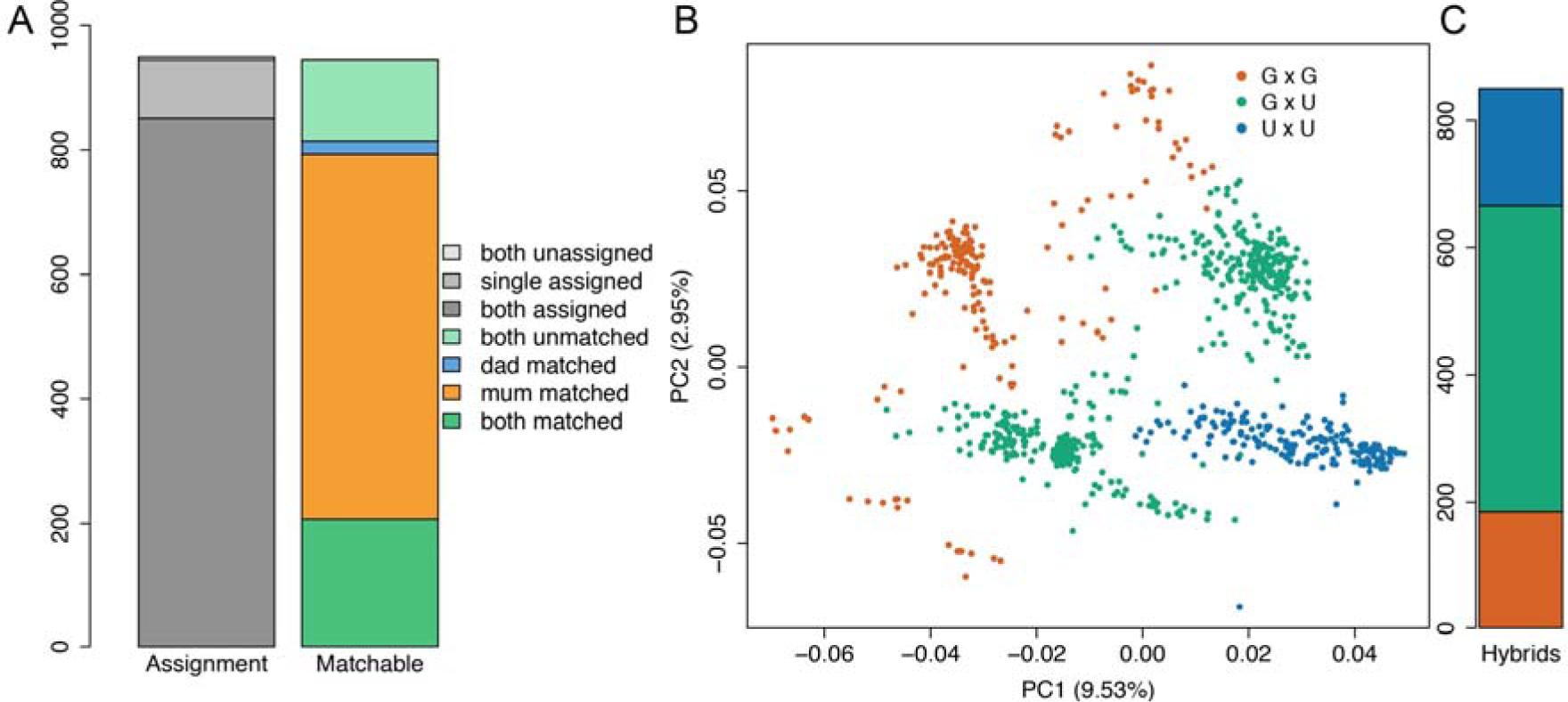
Summary of the parentage assignment and genetic structure. (A) Stacked bar plots from left to right represent the situations of parental assignment and matching, respectively. (B) First two principal components of a PCA test revealing population structure. Dots represent *E. urophylla* × *E.grandis* (green), *E. grandis* × *E. grandis* (dark orange), and *E. urophylla* × *E. urophylla* (dark blue) from the results of parentage assignment. (C) the number of each cross.

The assigned parent-offspring relationships largely agreed with the membership coefficients obtained from the genetic structure analysis (principal component analysis, PCA), reaffirming that the population consists of three types of crosses, two intra‐ and one inter-specific, namely, *E. grandis* × *E. grandis*, *E. urophylla* × *E. grandis* and *E. urophylla* × *E. urophylla* (Figure 1B). In contrast, the registered pedigree stated that all individuals were derived from *E. urophylla* × *E. grandis* crosses. For the 850 offspring where both parents could be assigned using SNP data, 489 (57.5%) were interspecific *E. grandis* x *E. urophylla* hybrids, 176 (20.7%) were intraspecific *E. grandis* and 185 (21.8%) were intraspecific *E. urophylla* (Figure 1C).

### 3.2. Estimates of variance components and heritability

Phenotypic data were adjusted by either removing spatial effects or by removing variation due to blocks in order to eliminate environmentally induced noise before fitting the additive and non-additive models. Height at age three years was adjusted with the use of spatial effects whereas other traits were adjusted for random block effects only since no autocorrelation was observed between rows and columns for these traits. Variance component and heritability estimates for all adjusted traits as well as AIC values for the nine different models (three genetic effect combinations with three relationship matrices) are presented (Table 2). Comparing A and AD models under the three relationship matrices, genomic-based models and pseudo-pedigree based models demonstrated very similar results in that the additive variance components estimated by the A models were much larger than those estimated by the AD models for growth traits. A large dominance variance was detected for these traits drawing variance from the additive one, suggesting that the additive and dominance variances are not independent. Greater additive variance components were detected for both genomic-based and pseudo-pedigree based models for wood traits. Dominance variance could only be found for basic density when using a genomic-based model and for pulp yield only when using a pseudo-pedigree based model. Results of models using the uncorrected registered-pedigree relationship matrix displayed a different and dramatic opposite trend with no evidence for dominance variance for growth traits while large dominance variances were detected for wood traits. For the ADE models we were not able to obtain results for the PADE and P_exp_ADE models due to matrix singularities that prevented the REML algorithm from converging. This probably occurs due to the shallow pedigree and that some variance components fall outside of the boundaries (zero or negative) that makes estimation impossible. We did detect epistatic variances for most growth traits under the GADE model, but no epistatic variance components were detected for wood traits.

**Figure 2.**
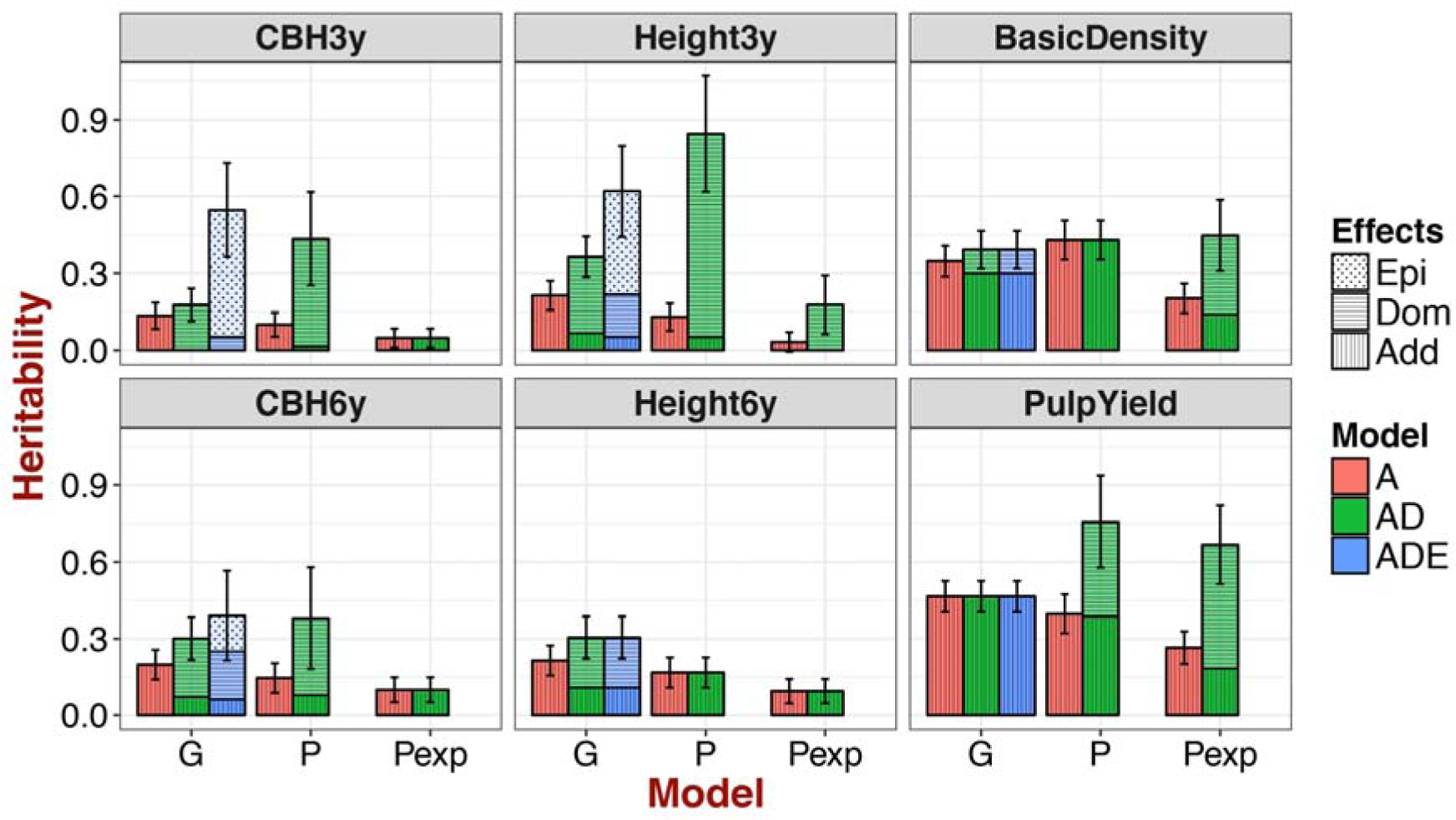
Narrow and broad sense heritability based on different models. Coloured boxes represent the different models used, where red indicate the additive model, green indicate the additive+dominance model and blue indicate the additive+dominance+epistasis model. Fill patterns represent different genetic effects, vertical lines denote additive effects, horizontal lines denote dominance effects and dots denote epistasis effects By combining both colour and fill patterns boxes, results from each model is displayed as separate specific genetic effects. The ADE model did not converge when we were using the pseudo-pedigree (P) and registered pedigree (P_exp_) to compute relationships among individuals for estimation. Black bars indicate the standard error of total genetic variance.

Narrow (h^2^) and broad-sense heritabilities (H^2^) were estimated for models using different relationship matrices (Figure 2). Generally, the additive effects decreased when non-additive effects were observed for AD and ADE models and large non-additive effects were obtained for growth traits where H^2^ increased more than 50%. In contrast, h^2^ of wood property traits were higher than growth traits and we also observed only slightly increases from h^2^ to H^2^ for these traits. Furthermore, standard errors (SE) of H^2^ were greater than SE for h^2^, but the SEs were generally smaller for genomic-based estimates compared to pedigree-based estimates for all traits.

**Table 2.**
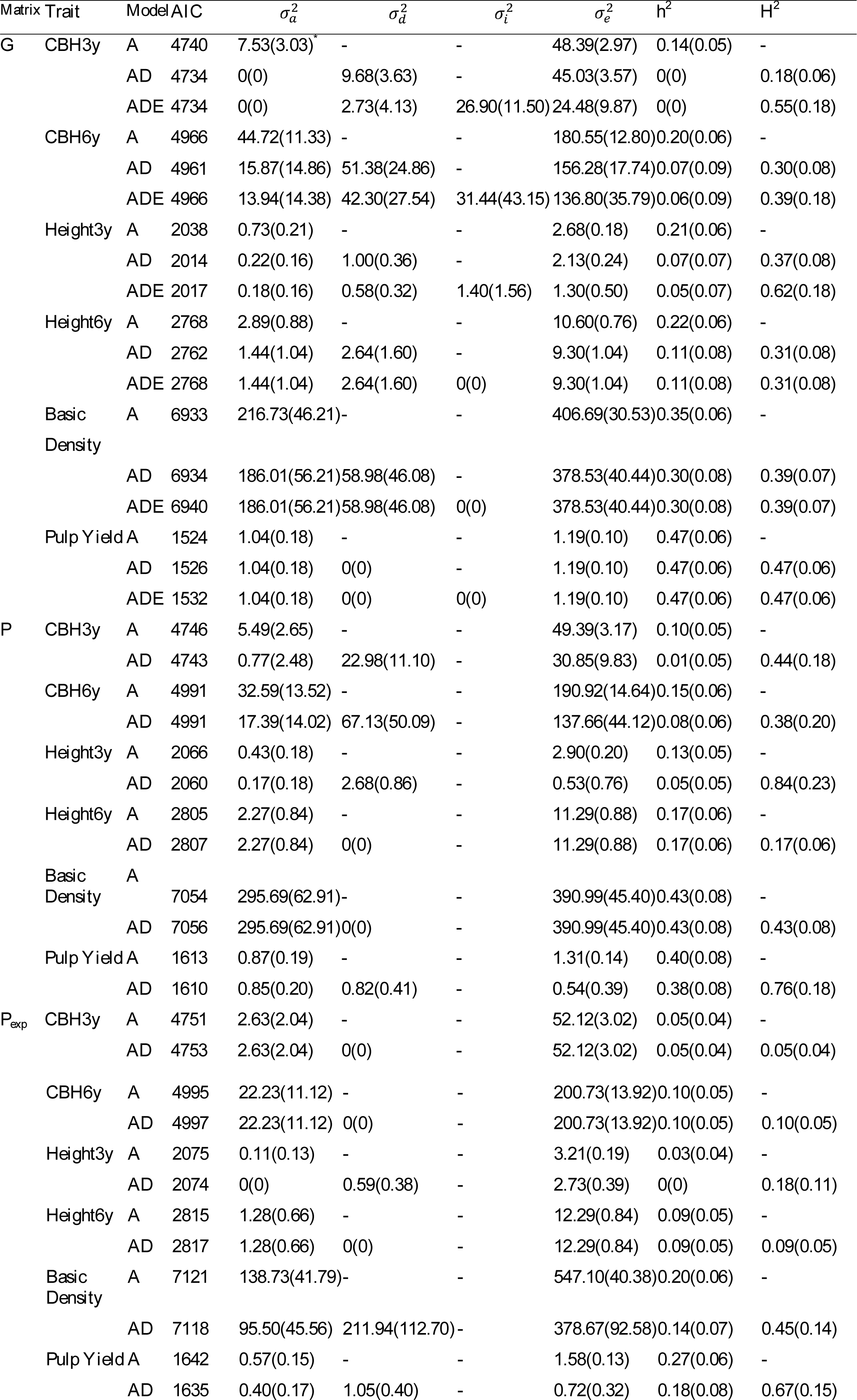
Summary of AIC, additive (
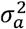
), dominance (
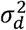
), epistasis (
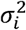
) and residual variances (
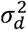
) and narrow− (h^2^) and broad-sense heritability (H^2^) of genetic models by accounting for genetic matrices

| Matrix | Trait | Model | AIC | $\sigma_a^2$ | $\sigma_d^2$ | $\sigma_i^2$ | $\sigma_e^2$ | $h^2$ | $H^2$ |
| --- | --- | --- | --- | --- | --- | --- | --- | --- | --- |
| G | CBH3y | A | 4740 | 7.53(3.03)* | - | - | 48.39(2.97) | 0.14(0.05) | - |
|  |  | AD | **4734** | 0(0) | 9.68(3.63) | - | 45.03(3.57) | 0(0) | 0.18(0.06) |
|  |  | ADE | 4734 | 0(0) | 2.73(4.13) | 26.90(11.50) | 24.48(9.87) | 0(0) | 0.55(0.18) |
|  | CBH6y | A | 4966 | 44.72(11.33) | - | - | 180.55(12.80) | 0.20(0.06) | - |
|  |  | AD | **4961** | 15.87(14.86) | 51.38(24.86) | - | 156.28(17.74) | 0.07(0.09) | 0.30(0.08) |
|  |  | ADE | 4966 | 13.94(14.38) | 42.30(27.54) | 31.44(43.15) | 136.80(35.79) | 0.06(0.09) | 0.39(0.18) |
|  | Height3y | A | 2038 | 0.73(0.21) | - | - | 2.68(0.18) | 0.21(0.06) | - |
|  |  | AD | **2014** | 0.22(0.16) | 1.00(0.36) | - | 2.13(0.24) | 0.07(0.07) | 0.37(0.08) |
|  |  | ADE | 2017 | 0.18(0.16) | 0.58(0.32) | 1.40(1.56) | 1.30(0.50) | 0.05(0.07) | 0.62(0.18) |
|  | Height6y | A | 2768 | 2.89(0.88) | - | - | 10.60(0.76) | 0.22(0.06) | - |
|  |  | AD | **2762** | 1.44(1.04) | 2.64(1.60) | - | 9.30(1.04) | 0.11(0.08) | 0.31(0.08) |
|  |  | ADE | 2768 | 1.44(1.04) | 2.64(1.60) | 0(0) | 9.30(1.04) | 0.11(0.08) | 0.31(0.08) |
|  | Basic Density | A | **6933** | 216.73(46.21) | - | - | 406.69(30.53) | 0.35(0.06) | - |
|  |  | AD | 6934 | 186.01(56.21) | 58.98(46.08) | - | 378.53(40.44) | 0.30(0.08) | 0.39(0.07) |
|  |  | ADE | 6940 | 186.01(56.21) | 58.98(46.08) | 0(0) | 378.53(40.44) | 0.30(0.08) | 0.39(0.07) |
|  | Pulp Yield | A | **1524** | 1.04(0.18) | - | - | 1.19(0.10) | 0.47(0.06) | - |
|  |  | AD | 1526 | 1.04(0.18) | 0(0) | - | 1.19(0.10) | 0.47(0.06) | 0.47(0.06) |
|  |  | ADE | 1532 | 1.04(0.18) | 0(0) | 0(0) | 1.19(0.10) | 0.47(0.06) | 0.47(0.06) |
| P | CBH3y | A | 4746 | 5.49(2.65) | - | - | 49.39(3.17) | 0.10(0.05) | - |
|  |  | AD | 4743 | 0.77(2.48) | 22.98(11.10) | - | 30.85(9.83) | 0.01(0.05) | 0.44(0.18) |
|  | CBH6y | A | 4991 | 32.59(13.52) | - | - | 190.92(14.64) | 0.15(0.06) | - |
|  |  | AD | 4991 | 17.39(14.02) | 67.13(50.09) | - | 137.66(44.12) | 0.08(0.06) | 0.38(0.20) |
|  | Height3y | A | 2066 | 0.43(0.18) | - | - | 2.90(0.20) | 0.13(0.05) | - |
|  |  | AD | 2060 | 0.17(0.18) | 2.68(0.86) | - | 0.53(0.76) | 0.05(0.05) | 0.84(0.23) |
|  | Height6y | A | 2805 | 2.27(0.84) | - | - | 11.29(0.88) | 0.17(0.06) | - |
|  |  | AD | 2807 | 2.27(0.84) | 0(0) | - | 11.29(0.88) | 0.17(0.06) | 0.17(0.06) |
|  | Basic Density | A | 7054 | 295.69(62.91) | - | - | 390.99(45.40) | 0.43(0.08) | - |
|  |  | AD | 7056 | 295.69(62.91) | 0(0) | - | 390.99(45.40) | 0.43(0.08) | 0.43(0.08) |
|  | Pulp Yield | A | 1613 | 0.87(0.19) | - | - | 1.31(0.14) | 0.40(0.08) | - |
|  |  | AD | 1610 | 0.85(0.20) | 0.82(0.41) | - | 0.54(0.39) | 0.38(0.08) | 0.76(0.18) |
| P<sub>exp</sub> | CBH3y | A | 4751 | 2.63(2.04) | - | - | 52.12(3.02) | 0.05(0.04) | - |
|  |  | AD | 4753 | 2.63(2.04) | 0(0) | - | 52.12(3.02) | 0.05(0.04) | 0.05(0.04) |

|  |  |  |  |  |  |  |  |  |
| --- | --- | --- | --- | --- | --- | --- | --- | --- |
| CBH6y | A | 4995 | 22.23(11.12) | - | - | 200.73(13.92) | 0.10(0.05) | - |
|  | AD | 4997 | 22.23(11.12) | 0(0) | - | 200.73(13.92) | 0.10(0.05) | 0.10(0.05) |
| Height3y | A | 2075 | 0.11(0.13) | - | - | 3.21(0.19) | 0.03(0.04) | - |
|  | AD | 2074 | 0(0) | 0.59(0.38) | - | 2.73(0.39) | 0(0) | 0.18(0.11) |
| Height6y | A | 2815 | 1.28(0.66) | - | - | 12.29(0.84) | 0.09(0.05) | - |
|  | AD | 2817 | 1.28(0.66) | 0(0) | - | 12.29(0.84) | 0.09(0.05) | 0.09(0.05) |
| Basic<br>Density | A | 7121 | 138.73(41.79) | - | - | 547.10(40.38) | 0.20(0.06) | - |
|  | AD | 7118 | 95.50(45.56) | 211.94(112.70) | - | 378.67(92.58) | 0.14(0.07) | 0.45(0.14) |
| Pulp Yield | A | 1642 | 0.57(0.15) | - | - | 1.58(0.13) | 0.27(0.06) | - |
|  | AD | 1635 | 0.40(0.17) | 1.05(0.40) | - | 0.72(0.32) | 0.18(0.08) | 0.67(0.15) |

\* Standard error (SE) is represented in parentheses.

* Standard error (SE) is represented in parentheses.

For all traits, the best model was obtained when using a genomic-based relationship matrix showing AIC values that were lower than for any of the other two relationship matrices. The GAD model was the best model for growth traits while the GA model was the best for wood traits (Table 2), which suggest that significant dominance effects can be detected for growth but not for wood traits whereas epistasis effects seemly play a minor role in all traits even though we can detect large epistatic variances for growth traits. We further studied the overall degree of dependency between the model variance estimates. We plotted the cumulative proportion of variance explained by the eigenvalues of the different models, relative to the diagonal representing an orthogonal correlation matrix (Figure S1). We found that the GAD outperformed the pedigree-based models (PAD and P_exp_AD) as indicated by closer adhering to the ideal scenario where the variance components are completely independent (diagonal line in Figure S1). Finally, since the GADE model does not have a corresponding model for the pedigree methods, GADE was plotted only against the diagonal line for reference (Figure S1).

### 3.3. Model fit and predictive ability

Model fit was estimated using the full data set (Table S1). The correlation between breeding values and phenotypes *r*(*Â_full_*,*Y_full_*) was only slightly lower for AD or ADE models compared to A only models for traits where we detected the contribution of non-additive variance. The correlations between genetic values and phenotypes *r*(*Ĝ_full_*,*Y_full_*) were higher than the corresponding correlations between phenotypes and breeding values *r*(*Â_full_*,*Y_full_*), with values varying between 0.8-0.95. With respect to the different relationship matrices that we used to fit models we found that the pseudo-pedigree based model in general had higher fit values than models using other relationship matrices. The registered-pedigree based model showed the lowest correlation for growth traits, whereas no marked differences were detected for wood traits (Table S1).

**Figure 3.**
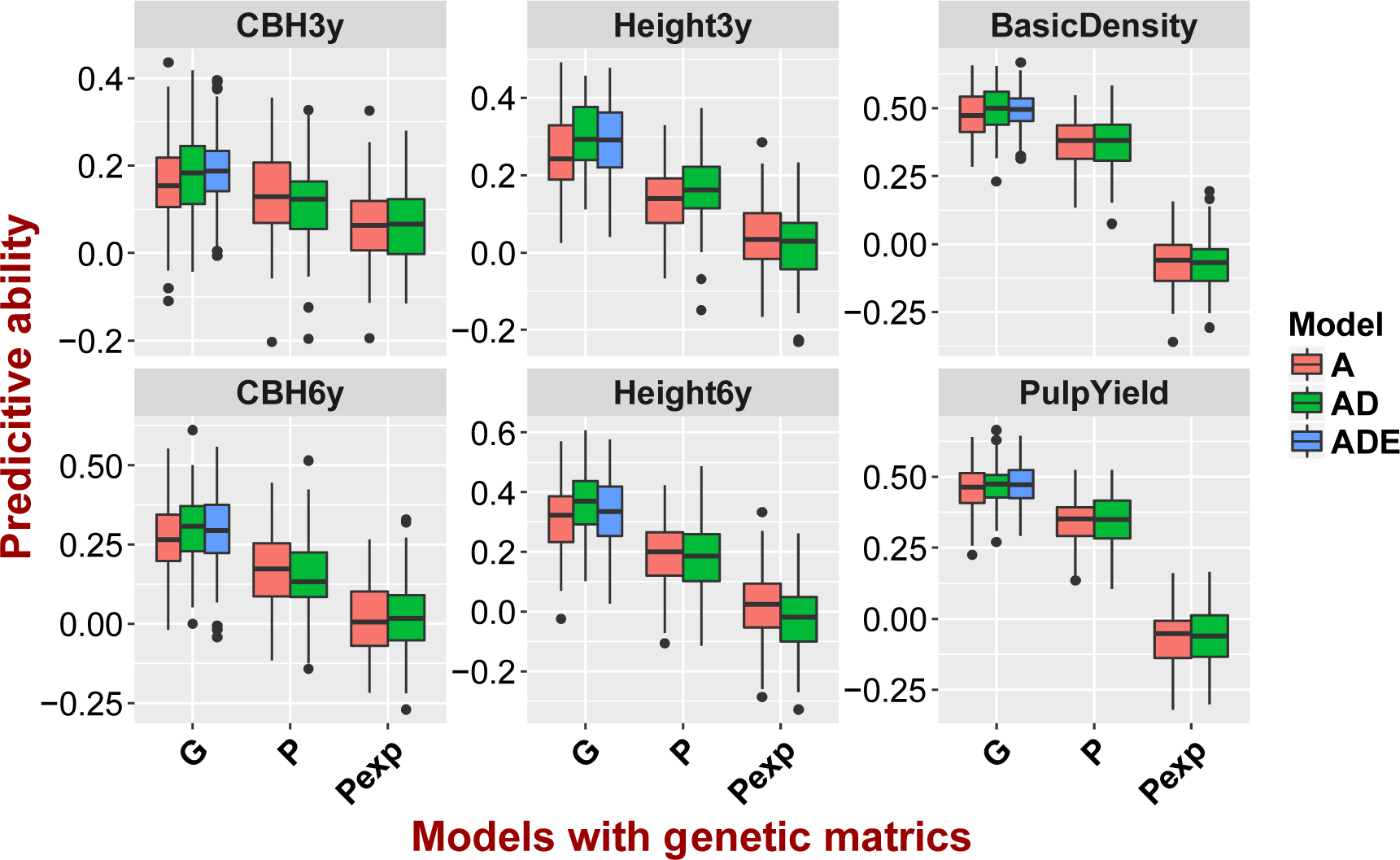
Predictive abilities for different models for each of the six traits. Boxplots showing the distribution of predictive ability over 100 replicates of ten-fold cross-validation from additive (A) (red), additive + dominance (AD) (green), and additive + dominance + epistatic (ADE) (blue) models estimated by genomic (G), pseudo-pedigree (P) and registered pedigree (P_exp_) based relationships.

Boxplots of the predictive ability of breeding values *r*(*Â_vali_*,*Y_vali_*) and genetic values *r*(*Ĝ_vali_*,*Y_vali_*) for the pedigree-based and marker-based models based on ten-fold cross-validation are shown in Figure 3. In general, and as expected, predictive abilities were lowest for the register-pedigree based models for all traits, ranging from −0.07 to 0.13. Furthermore, for genomic-based models (GA, GAD and GADE), a slight decrease in the predictive abilities of breeding values were observed (ranging from 0.14 to 0.31 across traits) when non-additive effects were included, while significantly higher predictive abilities were obtained for total genetic value (ranging from 0.19 to 0.36 across traits) when compared to breeding value for growth traits (Table 3). Overall, higher predictive abilities were observed for wood traits (0.5 for basic density and 0.44 for pulp yield) but there were no difference between predictive abilities for breeding value and total genetic value for these traits.

**Table 3.** The mean of predictive ability of breeding and genetic values for genetic models by accounting for genetic matrices

| Trait | Model | Matrix |  |  |  |  |  |
| --- | --- | --- | --- | --- | --- | --- | --- |
| | | G | | P | | $P_{exp}$ | |
| | | $r(\hat{A}_{vali}, \hat{Y}_{vali})^*$ | $r(\hat{G}_{vali}, \hat{Y}_{vali})^{**}$ | $r(\hat{A}_{vali}, \hat{Y}_{vali})$ | $r(\hat{G}_{vali}, \hat{Y}_{vali})$ | $r(\hat{A}_{vali}, \hat{Y}_{vali})$ | $r(\hat{G}_{vali}, \hat{Y}_{vali})$ |
| CBH3y | A | 0.16(0.10)*** | - | 0.13(0.09) | - | 0.06(0.09) | - |
|  | AD | 0.14(0.10) | 0.18(0.10) | 0.12(0.09) | 0.14(0.09) | 0.07(0.09) | 0.06(0.09) |
|  | ADE | 0.15(0.08) | **0.19(0.08)** | - | - | - | - |
| CBH6y | A | 0.27(0.11) | - | 0.18(0.12) | - | 0.01(0.12) | - |
|  | AD | 0.26(0.11) | **0.3(0.11)** | 0.14(0.11) | 0.15(0.12) | 0.01 (0.11) | 0.02(0.11) |
|  | ADE | 0.22(0.11) | 0.29(0.12) | - | - | - | - |
| Height3y | A | 0.26(0.10) | - | 0.13 (0.09) | - | 0.04(0.09) | - |
|  | AD | 0.23(0.10) | **0.30(0.08)** | 0.10(0.09) | 0.17(0.09) | 0.02(0.10) | 0.02(0.10) |
|  | ADE | 0.24(0.10) | 0.29(0.10) | - | - | - | - |
| Height6y | A | 0.32(0.11) | - | 0.19(0.11) | - | 0.02(0.12) | - |
|  | AD | 0.31(0.12) | **0.36(0.11)** | 0.19(0.12) | 0.19(0.12) | 0.01(0.11) | -0.02(0.12) |
|  | ADE | 0.3(0.13) | 0.33(0.12) | - | - | - | - |
| Basic Density | A | 0.47(0.08) | - | 0.37(0.09) | - | -0.07(0.10) | - |
|  | AD | 0.48(0.08) | **0.5(0.07)** | 0.38(0.10) | 0.38(0.10) | -0.06(0.10) | -0.07(0.10) |
|  | ADE | 0.48(0.07) | 0.49(0.07) | - | - | - | - |
| Pulp Yield | A | **0.46(0.08)** | - | 0.34(0.08) | - | -0.07(0.10) | - |
|  | AD | 0.45(0.08) | 0.46(0.08) | 0.34(0.09) | 0.34(0.09) | -0.06(0.10) | -0.06(0.10) |
|  | ADE | 0.44(0.08) | 0.46(0.08) | - | - | - | - |

\*\*\* Standard error (SE) is represented in parentheses.

* Correlation between phenotypes and breeding values on validation data set;

** Correlation between phenotypes and genetic values on validation data set;

*** Standard error (SE) is represented in parentheses.

## 4. Discussion

In our study we used a mostly F_1_ hybrid population derived from crosses between two *Eucalyptus* species to estimate the relative importance of additive and non-additive effects for growth and wood quality traits using genomic-based and pedigree-based models. We also analysed the contribution of non-additive effects to the accuracy of genetic values prediction with models that assume different genetic relationship matrices and for traits with different genetic architectures. Estimates of dominance and epistatic variances for genomic-based models indicated that non-additive genetic effects had substantial contributions to total genetic variation of growth traits (CBH and height at ages three and six years). The models including non-additive genetic effects also predicted genetic values more accurately, compared to a model without non-additive genetic effects. We were also able to estimate epistatic variance using the genomic-based model for the single generation of full-sib families that was not possible using a pedigree model.

### 4.1. Non-additive effects have substantial contributions to the genetic variance in growth

Although additive effects play a major role in most traits, non-additive effects should not be neglected. Our results demonstrated considerable contributions of non-additive variance captured by SNPs to the phenotypic variance of growth traits. The dominance effects contributed a further 4-15% to the total phenotypic variance (Table 2). Our results are consistent with those reported by Bouvet et al [41] and Muñoz et al [29], where significant effects of dominance were seen for height in *Eucalyptus* and loblolly pine, respectively. Moreover, our study found that between 0 to 30% of the phenotypic variance could be attributed to epistatic variation depending on the age when measurements were taken. These results corroborate previous results in *Eucalyptus* [1, 11, 19, 21, 41, 59], and further stress the importance of taking non-additive effects into account when breeding *Eucalyptus* F_1_ hybrids for growth. On the other hand, only a slight dominance variance was observed for basic density and none for pulp yield and epistatic variance estimates were zero for both wood traits (Figure 2). These results are in line with findings from previous pedigree-based studies in pines [60, 61] and *E. globulus* [62], but contrasts with results using half-sib families with marker-based genetic models in white spruce, where a very high proportion of epistatic variance in wood density was reported [42]. Therefore, these results suggest that the contribution of non-additive effects, especially epistatic effects, are both trait, species and possibly germplasm specific.

Our results show that the inclusion of dominance effects reduced the estimated narrow-sense heritability by 50%-70% for growth traits. Narrow-sense heritabilities for growth traits were further decreased by 70%-90% when both dominance and epistasis were taken into account (Figure 2). This trend is expected from a theoretical standpoint [63] as a substantial proportion of the non-additive variances can be manifested as additive variance in an additive-only model depending on the distribution of allele frequencies. This phenomenon has also been confirmed experimentally in other studies [29, 30, 41]. Moreover, the narrow-sense heritability for growth traits in our study population are rather low, only about 0.2 (Table 2). The low heritability we observe is likely caused by the selection of superior trees prior to genotyping. Trees were selected based on their growth and that likely have reduced variation in growth traits (CBH and height) which is reflected in the low heritability estimates. Such prior selection is of course not optimal for evaluating genomics based breeding methods, since it reduces the standing genetic variation but likely represents a common decision in operational breeding programs where high genotyping costs limits genotyping to a subset of the available offspring.

### 4.2. Models including dominance effects slightly improve prediction accuracy for growth

We evaluated how the inclusion of non-additive genetic effects impacted the prediction ability. For genetic values, the prediction ability slightly increased when going from GA to GAD models, whereas we observed no significant increase or sometimes even slightly decrease of prediction ability when going from GAD to GADE models (Figure 3 and Table 3). This result indicates that adding dominance effect to the model can improve predictive ability for traits where considerable dominance variance is detected, which support empirical results in both plants [64] and animals [65, 66].

However, although a large proportion of non-additive genetic variances were observed in GAD and GADE models for growth traits, we only observe a relatively small improvement (roughly 10%) in predictive ability (Table 3). Moreover, in the pedigree-based models, including dominance effects did not improve and sometimes even reduced the prediction ability (Figure 3). The results are accompanied by large standard errors on the non-additive variances components estimated with the ratios of dominance variances to standard errors are 0.5-0.9 for genomic-based models. Estimates for epistatic variances are even worse with ratios all exceeding 1. Furthermore, standard errors of pedigree-based models were 130-200% larger than those obtained for the genomic-based methods (Table 2). Large standard errors suggest a higher level of confounding effects in the analysis and thus a reduced power to predict genetic values [56]. Looking deeper into the characteristics of study population, the 949 F1 progeny represents a rather large effective population size (72 *E.grandis* and 73 *E.urophylla* parents), the number of individual per family is often too small (median family size is 2.44) and 25% of the families are represented by a single individual. Such imbalance between families reduce our ability to decompose observed variances into causal variance components which in turn yields large standard errors. Again, the situation is even worse for estimation of epistatic effects. Simulation results suggest that including non-additive effects should improve prediction ability in situations when the population size is large, when families are equally represented and when models are updated across selection cycles to reassess the relationship between markers and QTLs [43]. In conclusion, we find that including dominance effects slightly improve prediction accuracy but only for genomic-based models.

### 4.3. Genomics-based models outperform pedigree-based counterparts

Not surprisingly, our study show that pseudo-pedigree based models are markedly better than models based on the originally uncorrected registered pedigree both for genetic variance components estimation and for prediction. Comparing these two pedigree based models, dominance variances were detected only for the PAD models for growth traits, and PA models captured much more additive variance than the P_exp_A models (Figure 2). More importantly, predictive ability was substantially improved by using the pseudo-pedigree based models instead of registered-pedigree models due to the large number of errors in the latter (Figure 3). These results indicated that parentage assignment using SNP data can be very helpful for correcting pedigrees and evidently capturing more genetic variance and increasing the accuracy of predicting breeding values/genetic values [67]. However, our results showed that the predictive ability was further improved by using the full genomic-based relationship matrices instead of the pseudo-pedigree based relationship matrices (Figure 3). One reason is that parentage assignment did not find parents for all offspring. More importantly, however, is the fact that the genomic-based relationship matrix provides the marked advantage of capturing both the Mendelian segregation term within full-sib families and the cryptic genetic links through unknown common ancestors, which are not available simply from pedigree data even if this is totally correct. This feature has been highlighted in previous genomic selection studies in forest trees (e.g. [41, 42]).

Our results also showed that standard errors of the estimates of dominance variance obtained with the pedigree-based models were larger than those obtained when employing genomic-based models, indicating that genetic markers have better ability to estimate dominance effects than using pedigrees. Vitezica et al. [56] used simulations to show that genomic models were more accurate to estimate variance components when compared to pedigree-based models as evidenced by the smaller standard errors estimated for genomic models. Misztal [68] reported that accurate pedigree-based estimation of dominance variance requires at least 20 times as much data as required for estimation of additive variance. Moreover, the pedigree-based models did not converge when epistatic effects were added whereas genomic-based model could successfully be used to estimate epistatic effects under shallow pedigree and without clonal tests. This result supports earlier studies showing that pedigree-based models are inadequate for separating additive and non-additive effects without clonal trials [27].

AIC values for the genomic-based relationship matrix model were significantly lower than those based on pedigree relationship matrices, further corroborating that genomic-based models outperform the pedigree-based counterparts (Table 2). In addition, when we compared pedigree‐ and genomic-based models using the cumulative proportion of variance explained by eigenvalues of the sampling variance–covariance matrix of variance component estimates, we found that for most traits where dominance variance was detected, the GAD model outperformed the PAD/PexpAD models, as the variance components for the GAD model are less confounded (i.e. cumulative lines closer to the diagonal line, Figure S1). This result also suggests that the genetic variance components are not typically completely independent of each other, in line with earlier studies [29, 69].

### 4.4. Implications for breeding

Tree breeding involves a long and difficult process including plus tree selection, grafting, controlled pollination, and field trials. Without strict control and proper labelling, any of these steps could result in pedigree errors with far-reaching negative impacts on the outcomes of a breeding program, including but not limited to over or underestimation of expected genetic gains from production forestry. We have shown that the availability of SNP data allows extensive correction of errors in the expected pedigree structure, and increased accuracy in estimating genetic variances and breeding values.

Including dominance effects in the prediction of traits controlled by loci with additive and dominance effects results in higher predictive ability for genotypic values. This will increase genetic gains for clonal selection and for the recurrent selection of superior mate pairs. As a proof-of concept, we compared the overlap among the top 100 performing individuals selected with the PA, PAD, GA and GAD models (Figure S2). For growth traits when comparing these four models, only~30-40% of the top 100 individuals were selected by all of them based on early age measurements at age three but the proportion increased to a quite acceptable level of 40-50% at harvest age of six years. This corroborates the critical importance of using growth data close to or preferably at harvest age to build genomic prediction models for optimal implementation of genomic selection for growth traits in *Eucalyptus*. For wood traits, however, more than 50% of the individuals overlapped, and up to 72 individuals were identified by all models for basic density. This result is particularly relevant because it shows great prospects to practice genomic selection already at the seedling stage for late expressing wood traits using SNP data.

Our predictive ability results also showed that using genomic realized relationships provides much improved prediction of complex phenotypes, both for breeding values and total genetic values, as more information is used. In addition, our study confirms that non-additive variation is prevalent in hybrid eucalypts for growth but not for wood quality traits. This realized-genetic based model by including non-additive effect has proven effective in animal breeding [70-72] and has also been advocated for plant breeding (reviewed in [73]). Such model can thus improve the efficiency and productivity of variety selection pipelines that are the most labour‐ and time-intensive component of a breeding cycle to arrive to elite planting material.

Accurate estimation of non-additive genetic variance using SNP data will also assist the choice of optimal tree breeding strategy, particularly for hybrid breeding programs. Simulation studies have shown that a synthetic breeding population composed by first or second generation hybrids might be the most cost effective in terms of gain per unit time for traits where there is less dominance variance and a positive correlation exists between performance of pure species and hybrids. However, for traits where gene action is primarily dominant, reciprocal recurrent selection with forward selection (RRS-SF) is probably the best breeding strategy [5]. Our results show an important contribution of dominance for growth but not for wood quality in the widely bred *E. grandis x E. urophylla* hybrid and would therefore require a compromise as far as the relative importance of wood basic density and pulp yield in the breeding objective, i.e. a linear combination of the traits of economic importance. While it remains to be seen whether dominance effects could also be expressed and satisfactorily captured in a synthetic breeding population, volume is typically a dominant trait in determining the benefits in short-rotation eucalypt [74], such that RRS-SF might still be the best option despite its much longer breeding cycle and logistic complexity. In any case, our work shows that the use of SNP data in breeding and the promising perspectives of adopting ultra-early genomic selection for all traits of economic importance in hybrid eucalypt will open new avenues to better evaluate the several options available to the breeder to optimize the breeding objective.

## Acknowledgements

We would like to thank Dr. Eduardo Pablo Cappa for his suggestions of data analysis. This research was conducted using the resources of the High Performance Computing Center North (HPC2N).

